# An explanation for the sister repulsion phenomenon in Patterson’s f-statistics

**DOI:** 10.1101/2024.02.17.580509

**Authors:** Gözde Atağ, Shamam Waldman, Shai Carmi, Mehmet Somel

## Abstract

Patterson’s f-statistics are among the most heavily utilized tools for analysing genome-wide allele frequency data for demographic inference. Beyond studying admixture, *f_3_* and *f_4_* statistics are also used for clustering populations to identify groups with similar histories. However, previous studies have noted an unexpected behaviour of f-statistics: multiple populations from a certain region systematically show higher genetic affinity to a more distant population than to their neighbours, a pattern that is mismatched with alternative measures of genetic similarity. We call this counter-intuitive pattern “sister repulsion”. We first present a novel instance of sister repulsion, where genomes from Bronze Age East Anatolian sites show higher affinity towards Bronze Age Greece rather than each other. This is observed both using *f_3_*- and *f_4_*-statistics, contrasts with archaeological/historical expectation, and also contradicts genetic affinity patterns captured using PCA or MDS on genetic distances. We then propose a simple demographic model to explain this pattern, where sister populations receive gene flow from a genetically distant source. We calculate *f_3_*- and *f_4_*-statistics using simulated genetic data with varying population genetic parameters, confirming that low-level gene flow from an external source into populations from one region can create sister repulsion in f-statistics. Unidirectional gene flow between the studied regions (without an external source) can likewise create repulsion. Meanwhile, similar to our empirical observations, MDS analyses of genetic distances still cluster sister populations together. Overall, our results highlight the impact of low-level admixture events when inferring demographic history using f-statistics.

## Introduction

Over the last decade, Patterson’s f-statistics (Patterson *et al*. 2012) have become one of the most commonly applied tools in population genetics studies involving the analysis of genome-wide data. Various f-statistics (*f_2_*, outgroup*-f_3_*, admixture*-f_3_*, and *f_4_*) quantify divergence, shared genetic drift and treeness among populations by comparing allele frequencies. These measurements enable the identification of population structure, differentiation and admixture events and help decipher the demographic history of the studied populations (Peter 2016; Harris and DeGiorgio 2017; Lipson 2020; Peter 2022). The description of f-statistics as being robust to heterogeneity of sample size/coverage and ascertainment biases (Patterson *et al*. 2012) has gained them high popularity.

The outgroup-*f_3_* statistic in the form of *f_3_(Outgroup; Pop2, Pop3)* measures shared drift between *Pop2* and *Pop3* relative to an outgroup, and its magnitude has been used as a measure of pairwise population affinity. The *f_4_*-statistic measures treeness, such that *f_4_(Outgroup, Pop1; Pop2, Pop3)* is expected to be 0 if *Pop1* shares the same amount of drift with *Pop2* and *Pop3*, i.e. *Pop1* is a real outgroup to the latter two populations. Significant deviation from 0 can be interpreted as evidence for admixture (Patterson *et al*. 2012), but the *f_4_*-statistic can also be used for studying population structure and shared demographic histories. If *Pop2* and *Pop3* share more of their demographic history than they do with *Pop1*, i.e. shared drift and/or similar admixture events that *Pop1* did not experience, and if there is no post-split admixture, we expect *f_4_(Outgroup, Pop1; Pop2, Pop3)* to be non-significant, while *f_4_(Outgroup, Pop2; Pop1, Pop3)* and *f_4_(Outgroup, Pop3; Pop1, Pop2)* to both be positive. Gene flow from *Pop1* into *Pop3* in the same model will also cause *f_4_(Outgroup, Pop1; Pop2, Pop3)* to be positive. Thus, in addition to testing admixture, reciprocal affinity in *f_4_*-tests has frequently been used as a tool for clustering populations based on allele frequency similarities, where clustering implies similar histories. The outgroup-*f_3_* statistic has similarly been used to evaluate shared history between populations.

Nevertheless, f-statistics are not always straightforward to interpret. Complications of diverse types and origins have been noted, such as possible misinterpretation of an outgroup as an admixture source when using admixture-*f_3_* (Patterson *et al*. 2012), or the sensitivity of *f_4_*-statistics to SNP ascertainment schemes as showcased for demographic modelling of African human populations using SNPs identified in non-African groups (Bergström *et al*. 2020; Flegontov *et al*. 2023). The most widespread type of issue arguably involves the interpretation of *f_4_* -statistics under the assumption of a simple demographic model where the real history is substantially more complicated. For instance, unrecognised genetic structure can be misinterpreted as admixture events (Durand *et al*. 2011; Eriksson and Manica 2012). A similar confounding effect can be created by backward gene flow, as shown for Neanderthal admixture estimates varying spuriously over time due to African gene flow into Eurasia (Petr *et al*. 2019a).

Previous work using f-statistics has noted another unexpected pattern that has yet remained largely unexplored, to our knowledge. Using *f_4_*-statistics, Kılınç et al. observed higher affinity of all Anatolian Neolithic groups to a Neolithic genome from Greece than to each other, a pattern that appeared difficult to interpret (Kılınç *et al*. 2017). Even more surprising was the observation that modern-day North African populations showed higher affinity to modern-day Sardinians than to each other using *f_3_-* and *f_4_*-statistics (Rodríguez-Varela *et al*. 2017). This pattern was hypothesized to stem from gene flow from sub-Saharan Africa into North Africa, but the case was not studied further.

Here, we provide another case of this pattern, which we call “sister repulsion”, by comparing genomes from the Bronze Age (BA) East Mediterranean. We then describe a simple theoretical model that can explain this effect, propose different scenarios involving external and regional gene flow to explain sister repulsion, and recapitulate the observed f-statistics using simulations under these models, with various conditions. Our results showcase how complex demographic histories can lead to counter-intuitive patterns of *f_3_-* and *f_4_-*statistics when used for studying shared population history.

## Results and Discussion

Our initial motivation was to investigate population structure and admixture patterns in the East Mediterranean during the Bronze Ages, a period which saw population growth and extensive interregional trade networks being established, as inferred from material culture data (Şahoğlu 2005; Kouka 2016; Massa 2016). We studied genetic affinities between published genomic data from n=6 BA populations from modern-day Greece (2400-1070 BCE) and n=5 populations from BA East Anatolia (3700-1303 BCE) (Lazaridis *et al*. 2017; Skourtanioti *et al*. 2020; Lazaridis *et al*. 2022b) (Table S1, Figure S2). We restricted the analysis to 1240K capture data to avoid confounding by technical factors (Margaryan *et al*. 2020) likely caused by reference bias (Rohland *et al*. 2022; Koptekin *et al*. 2023). We note that the results described henceforth are also observed using Central and West Anatolian BA populations, which we do not show here; we instead focus on East Anatolia as analyses involving populations from this region are the most counter-intuitive.

First, we explored genetic clustering among populations with Principal Components Analysis (PCA) and Multidimensional Scaling (MDS) analysis. In the PCA, where the ancient genomes are projected on a PC space calculated from modern-day West Eurasians, genomes from modern-day Greece cluster separately from those from East Anatolia (Figure S1). Then, we calculated the genetic distance between all pairs of populations using 1 - *f_3_(Yoruba; Pop1, Pop2)* to represent the distance between populations *Pop1* and *Pop2*. Applying MDS analysis on the resulting pairwise distance matrix (Fu *et al*. 2016), the populations showed a tendency to cluster by region in MDS space, i.e. Greece and Anatolia separated along Coordinate 1 (Figure 1A,B).

**Figure 1:**
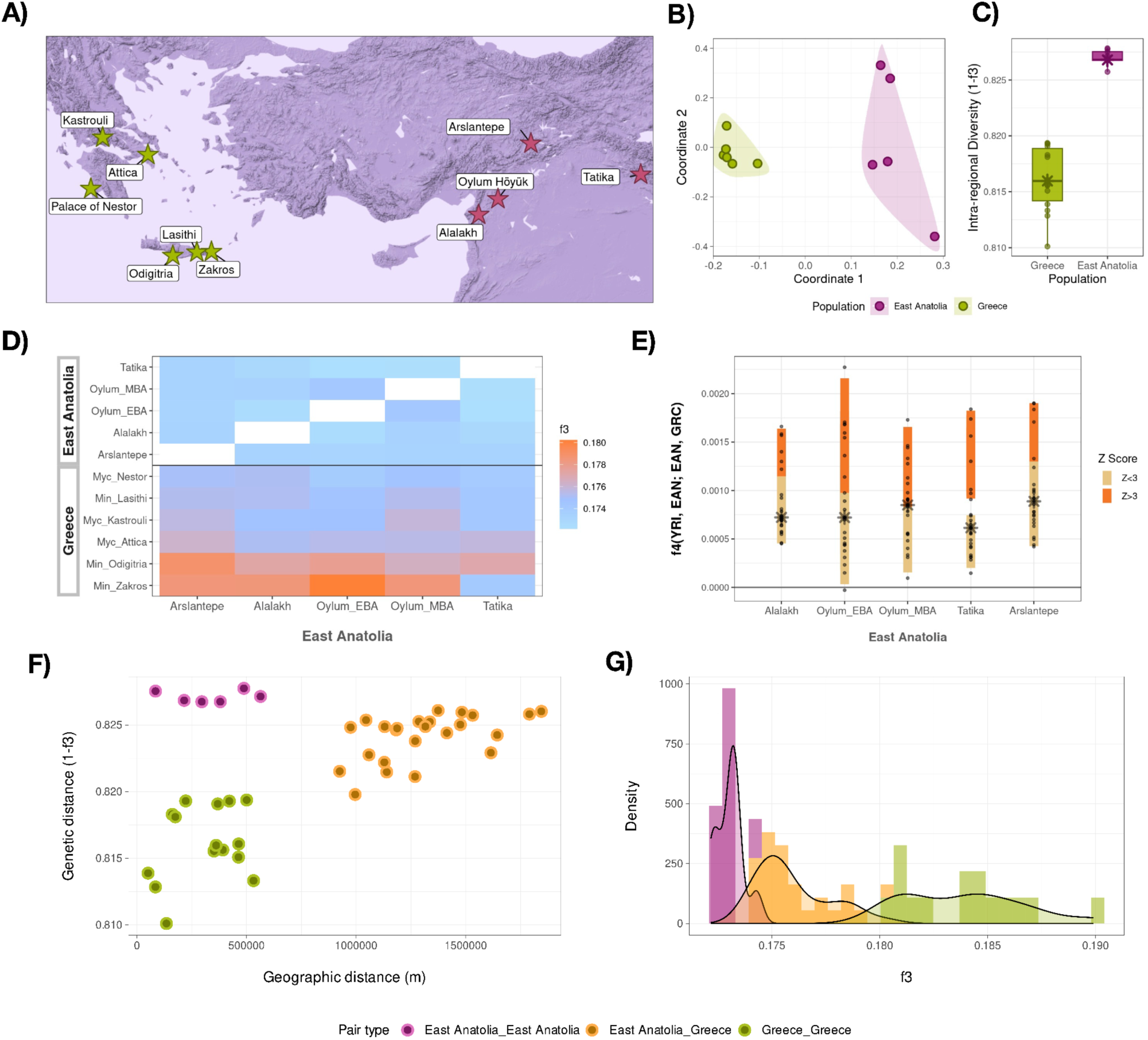
**(A)** Geographical distribution of the studied Bronze Age genomes from modern-day Greece and East Anatolia. **(B)** Genetic clustering of the populations in line with geography revealed by multidimensional scaling analysis (MDS), using 1 - outgroup- *f_3_* values as genetic distances. Figure S1 also presents a PCA, which also shows the absence of visible structure in these gene pools. **(C)** Intra-regional diversity of the populations measured as pairwise 1 - outgroup-*f_3_* values within regions. Each point represents comparisons of settlements from the same region. **(D)** *f_3_* statistics of the form *f_3_(Yoruba; East Anatolia, X)* where X corresponds to populations from modern-day Greece and East Anatolia. **(E)** *f_4_* statistics of the form *f_4_(Yoruba, East Anatolia; East Anatolia, Greece)*, where positive values depict higher affinity of BA East Anatolian populations towards contemporary populations from modern-day Greece. **(F)** Geographic versus genetic distances (1-f_3_) between and within regions. **(G)** Distribution of *f_3_* values between and within regions.

Intriguingly, however, *f_3_* statistics revealed a counter-intuitive trend. In contrast to the patterns observed in PCA and MDS, the values *f_3_(Yoruba; EAnatoliaY, GreeceX)* were consistently higher than the values *f_3_(Yoruba; EAnatoliaY, EAnatoliaZ)*, where *GreeceX* is one of the six populations from modern-day Greece, while *EAnatoliaY* and *EAnatoliaZ* are any of the five populations from East Anatolia (Figure 1B-E,G, Table S2). At face value, this would imply more similar histories between any East Anatolian population with those from modern-day Greece than other neighbouring East Anatolian populations. Plotting the genetic versus geographic distances, we observed a prominent lack of relationship within the East Anatolian populations (Figure 1F).

We next calculated *f_4_*-statistics of the form *f_4_(Yoruba, AnatoliaY; AnatoliaZ, GreeceX)*. In 99% of 120 comparisons we observed positive values (42% significant at Z>3, without correcting for multiple testing) (Figure 1D, Table S3). No *f_4_-*statistic was significantly negative, i.e. no East Anatolian group significantly shared more drift with one of its neighbours over a population from modern-day Greece. This starkly contrasts with the expected geographical and genetic clustering [given multiple observations of spatial population structure in ancient humans (e.g. Altınışık, Kazancı, *et al*. 2022; Antonio *et al*. 2024)], while it does mirror the outgroup-*f_3_* results.

We then asked whether population or time structure within the two regional groups could be behind this pattern, as the sampled individuals span several thousands of years. We investigated this using *f_4_*-statistics within one site in East Anatolia, Alalakh. We randomly split the n=25 genomes from Alalakh into two groups and calculated *f_4_(Yoruba, Group1; Group2, Greece)*, ten times independently. We observed positive *f_4_* values, or sister repulsion, in all ten Alalakh replicates (all *f_4_* values were significant except for one trial) (Table S4).

Assuming this is not caused by technical artifacts, such as contamination, there could be two explanations for this sister repulsion pattern. One may be gene flow from modern-day Greece into different East Anatolian populations or the ancestors of all East Anatolian populations. An alternative explanation and a possibly more likely scenario could involve admixture from an external and diverse source impacting East Anatolia. This would be similar to that hypothesized by Rodríguez-Varela and colleagues (2017) and analogous to the “outgroup case” described by Patterson and colleagues (2012).

The ancestral components of both Anatolia and Greek Early/Middle Bronze Age genomes comprise Neolithic Anatolia and eastern-related ancestry from Caucasus/Mesopotamia (Clemente *et al*. 2021; Lazaridis *et al*. 2022b; Skourtanioti *et al*. 2023). We hypothesized that additional external gene flow into East Anatolia (e.g. from the Levant, Zagros, or Mesopotamia) that BA populations from Greece did not receive could lead to differentiation within the former region and might explain the observed sister repulsion pattern. As a first test of this idea, we calculated intra-regional diversity within each site, using the pairwise statistic 1 - outgroup-*f_3_* among genomes within a site as the diversity measure. This revealed significantly higher median diversity in BA East Anatolia than in BA Greece (Wilcoxon rank sum test p=3e-09, Figure 1C, S3).

This supports the hypothesis that gene flow from a divergent source (so-called “deep ancestry”) might explain this pattern. We investigated the idea further using the scenario described in Figure 2. We define A and B as two sister populations, X as a more distant lineage, Z as the source of external gene flow into A and B, and O as outgroup to all. Here, A, B and X represent our focal populations; A and B could represent two BA East Anatolian groups, X could be a BA population from the region of modern-day Greece, and Z would be an unknown, distant source that admixes only with East Anatolia. The ancestors of multiple populations are denoted by their merged names (i.e. the ancestor population of A and B is called AB). Note that in the model, A and B are sisters; their demographic histories are the same except for receiving independent gene flow from Z.

**Figure 2:**
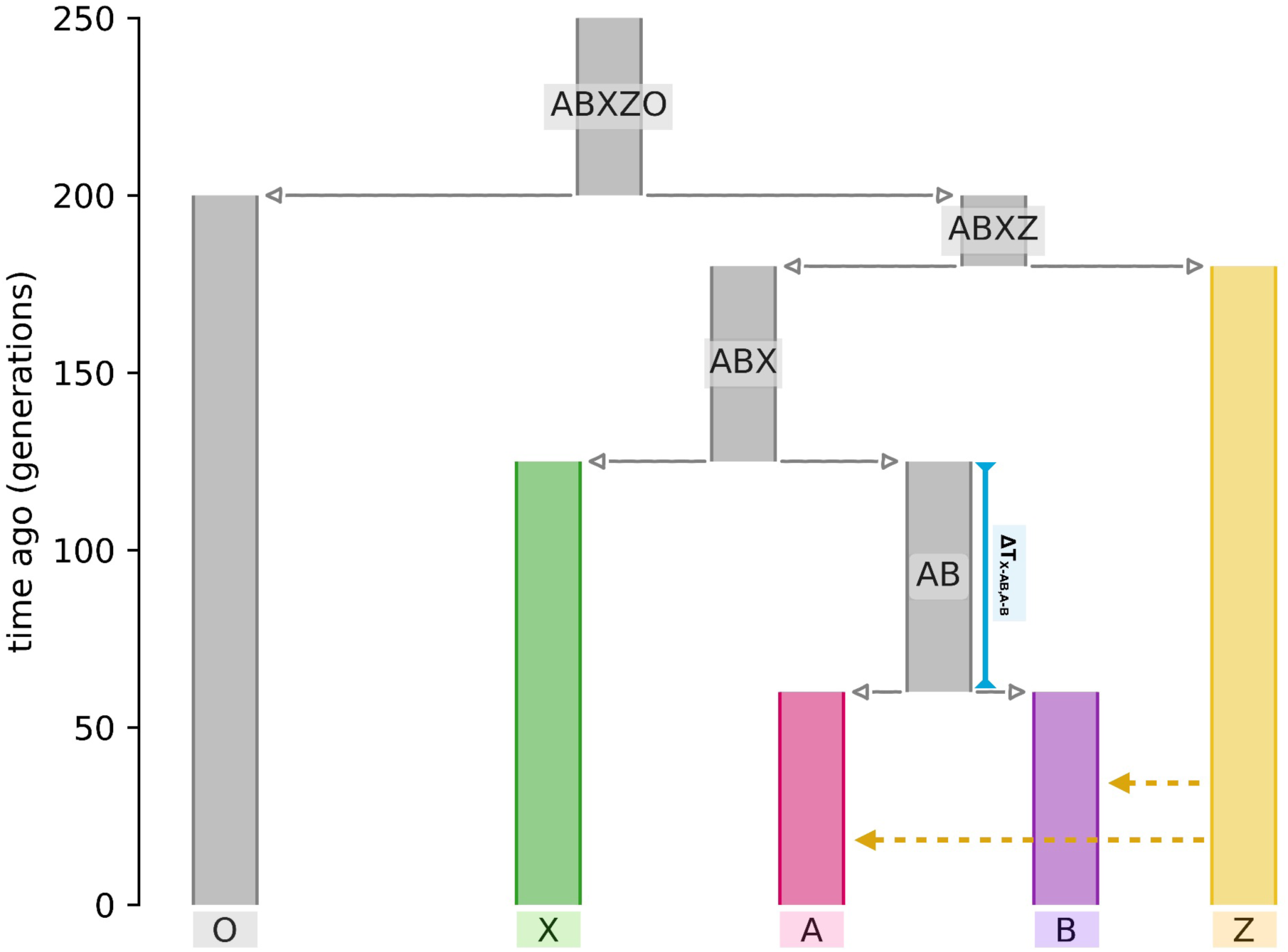
Demographic model used for the population genetic simulations. The ancestral populations are named as combinations of the descending population names. Population O is the outgroup used in the calculation of f-statistics. Dashed lines represent equal rates of continuous migration from Z to A and B per generation after the split between A and B and the present day. The blue line corresponds to the parameter ΔT_X-AB,A-B_.

We studied the theoretical conditions for the expected value of *F_4_(O, A; B, X)* to be positive in this model (we follow Patterson *et al*. in using upper case letters for expected values). The *F_4_* statistic can be decomposed into the sum of overlapping dirft paths between O to A and B to X. Under gene flow from Z into A and/or B, there emerge four alternative drift paths, described in Figure 3. These depend on *α*_A_ and *α*_B_, the probabilities of gene flow (admixture proportions) from Z into A and B. The expected *F_4_* is the weighted sum of the values of these paths. One of these paths produces a positive value, i.e. sister repulsion, which depends on the branch length *g* separating Z from the common ancestor of A, B, and X (branch ABX in Figure 2), on the branch length *e* separating X from the common ancestor of A and B (branch AB in Figure 2) and also on the admixture proportions *α*_A_ and *α*_B_. Note that the branch lengths represent drift and will be a function of time and N_e_.

**Figure 3:**
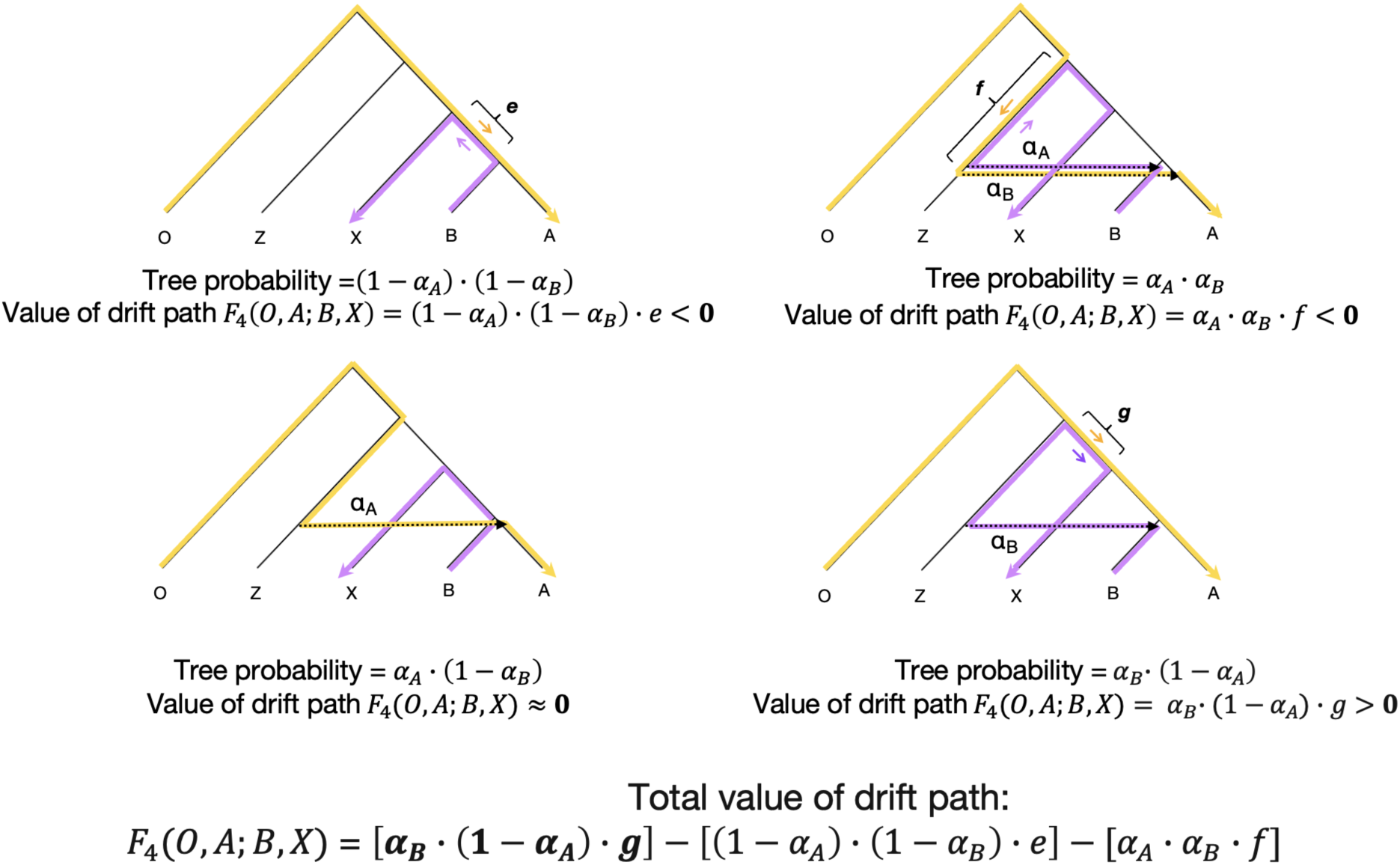
Expected *F_4_* values under distant admixture. The four trees with arrows show four possible drift paths that can contribute to *F_4_*(O, A; B, X). Here, sister populations A or B have independently received gene flow (dashed arrow) from the distant branch Z with admixture proportions ɑ_A_ and ɑ_B_, respectively, after their split from each other. The probability of each path and its value as a product of its probability and the drift magnitude (indicated by letters *e*, *f*, and *g*) are shown below each tree. The expected average *F_4_* value (Patterson et al. 2012) is shown at the bottom, with the positive component in bold. The small arrows indicate the direction of the drift paths from O->A and B->X, which determine the sign of that *F_4_* path.

Now let us we assume that the admixture proportions in A and B are the same (*α*_A_ = *α*_B_) and small (so that *α*^2^ is negligible). Then, *g* ⋅ *α* > *e* would fulfill the condition of a positive *F_4_*, i.e. sister repulsion. For instance, in a simplistic scenario where all populations had the same N_e_ and the admixture proportions *α* were the same, if BA populations from the regions of modern-day Greece and E Anatolia had split 30 generations ago (∼900 years), the latter received gene flow from a population that had split 2000 generations ago (∼60,000 years), an admixture proportion of >0.015 would suffice to create a positive *F_4_* value.

Having shown sister repulsion is possible theoretically, we next studied the behaviour of f-statistics numerically with population genetic simulations. Using the general model in Figure 2 we ran multiple simulations where we varied the parameters m_Z->A.B_, Ne_Z_, Ne_AB_, Ne_ABX_, T_A-B_, and T_X-AB_ (Table 1). The continuous migration m_Z->A.B_ was set to start subsequent to the A-B split. For simplicity, we refer to T_A-B_ and T_X-AB_ as a single variable that shows T_X-AB_ -T_A-B_, which we call ΔT_X-AB,A-B_. In each case we sampled 25 genomes each from O, X, A, and B, and calculated various outgroup-*f_3_* and *f_4_*-statistics.

**Table 1:** Parameter values used for the main population genetic simulations. Migration rates are given per generation, divergence times are given in generations.

| Parameter | Values | Definition |
| --- | --- | --- |
| $m_{Z \rightarrow A,B}$ | 0, 0.001, 0.005, 0.01, 0.1 | Migration rate from Z to A and B |
| $Ne_Z$ | 5k, 15k, 35k | Effective population size of source Z |
| $Ne_{AB}$ | 5k, 15k, 35k | Effective population size of ancestral AB |
| $Ne_{ABX}$ | 1k, 3k, 5k | Effective population size of ancestral ABX |
| $T_{A-B}$ | 30, 60, 90 | Divergence time of A and B |
| $T_{X-AB}$ | 100, 125, 150 | Divergence time of X and AB |
| $\Delta T_{X-AB,A-B}$ | 10, 65, 120 | Difference between $T_{A-B}$ and $T_{X-AB}$ |
| $Ne_X^*$ | 1k, 3k, 5k | Effective population size of X |
| $m_{X \rightarrow A,B}^*$ | 0, 0.001, 0.005, 0.01, 0.1 | Migration rate from X to A and B |

\* The given parameter values for $Ne_X$ and $m_{X \rightarrow A,B}$ were tested only for scenarios where $Ne_{ABX} = 1k$ and $\Delta T_{X-AB,A-B} = 65$ . In the remaining scenarios, $Ne_X$ was fixed at 5k and $m_{X \rightarrow A,B}$ at 0.

In this model (Figure 2) one would naively anticipate *f_4_(O, A; B, X)* values to be negative since the populations A and B appear to share more similar histories with each other than with X, as in tree A in Figure 3. Indeed, in the absence of gene flow from Z into A/B (m=0), *f_4_* statistics are mostly negative, reflecting higher shared drift between A and B than with X (Figure 4, Figure S2). However, *f_4_* values change as we allow migration from Z into A/B (from m=0 to m=0.1), although in non-linear fashion. When we increased migration from 0 to 0.01, we observed a monotonic rise in *f_4_(O, A; B, X)* values, i.e. an increasing trend towards sister repulsion (Figure 4). Under specific conditions, e.g. Ne_ABX_ = 1k, 0.001≤m<0.1 and ΔT_X-AB,A-B_ =10, all *f_4_(O, A; B, X)* values are positive (Figure 4, S4). The same applies to *f_4_(O, B; A, X)*. This sister repulsion pattern is analogous to that observed in the empirical data where two East Anatolian populations showed higher affinity to populations from modern-day Greece than to each other (Figure 1D). We also repeated our random split approach explained above for the real data in a simulated scenario for individuals in population A and again obtained all positive *f_4_* values (Table S5). In summary, we can replicate the empirical result effectively under specific conditions.

**Figure 4:**
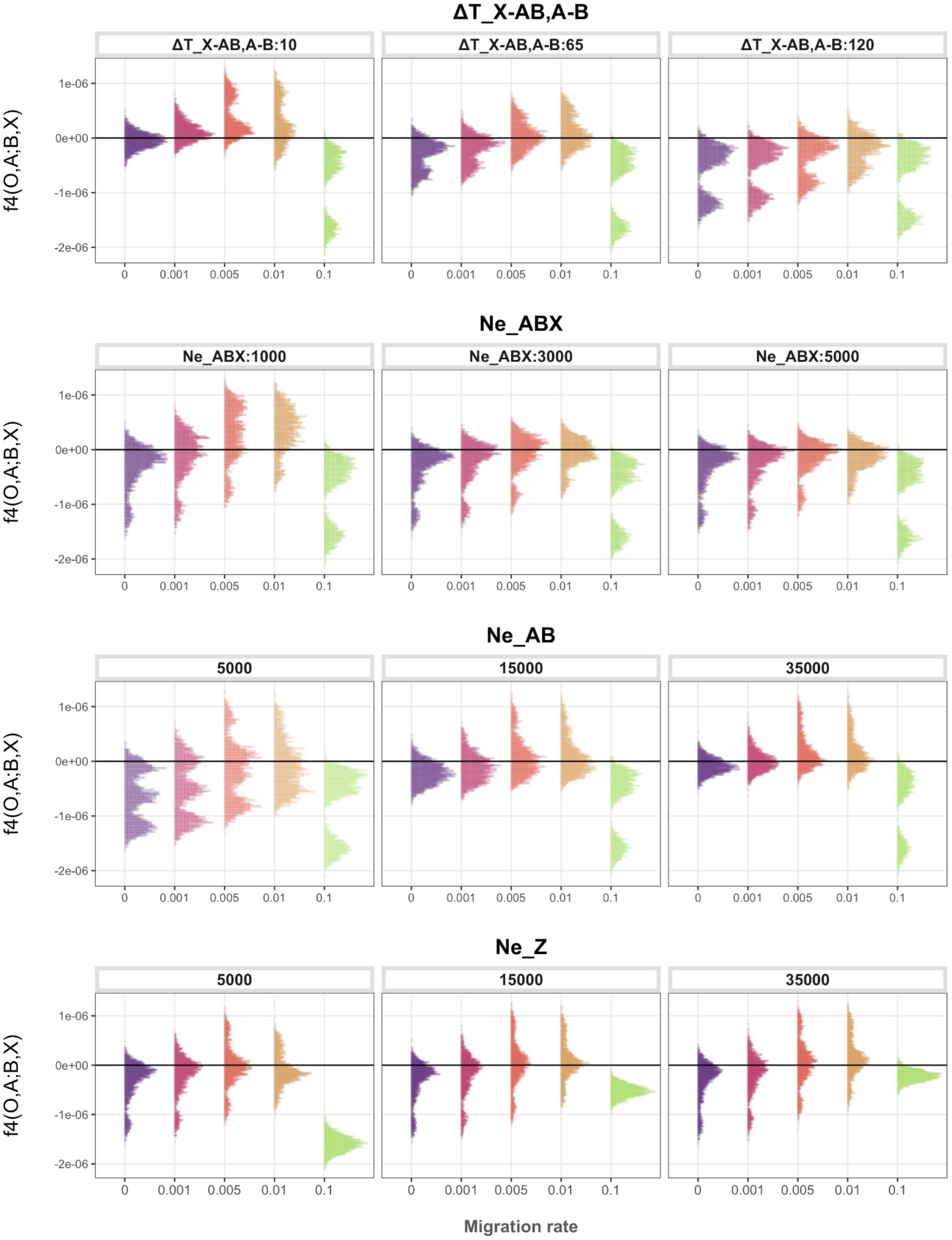
Change in *f_4_* statistics of the form *f_4_(O, A; B, X)* depending on the varying parameters in population genetic simulations, where negative values represent higher affinity between A and B, and positive values higher affinity between A and X. The panels for each parameter contain all values of the remaining studied parameters and are explained in Table 1. The multimodal distribution shapes arise because each distribution is collected from a range of simulations run under a range of parameters with diverse effects on the *f_4_* statistic.

We note several conditions driving more positive *f_4_(O, A; B, X)* values under low-level migration in our model (Figures 4,5; Table 1).

- Low ΔT_X-AB,A-B_ (Figure 4 first row): this means A and B share less drift relative to X (the phylogeny is more star-like). Also, because T_AB-X_ is fixed in the model, low ΔT_X-AB,A-B_ leads to T_A-B_ going back in time (Figure 2) and therefore A and B each being impacted by Z-sourced continuous migration for additional generations (i.e. larger *α* in Figure 3).
- Low Ne_ABX_: this causes higher shared drift between A, B and X (i.e. larger *g* in Figure 3).
- High Ne_AB_: this leads to lower shared drift between A and B (i.e. smaller *e* in Figure 3).
- High Ne_Z_: this causes Z to be subject to less drift, such that the negative branch length *f* in Figure 3 becomes smaller in magnitude.

**Figure 5:**
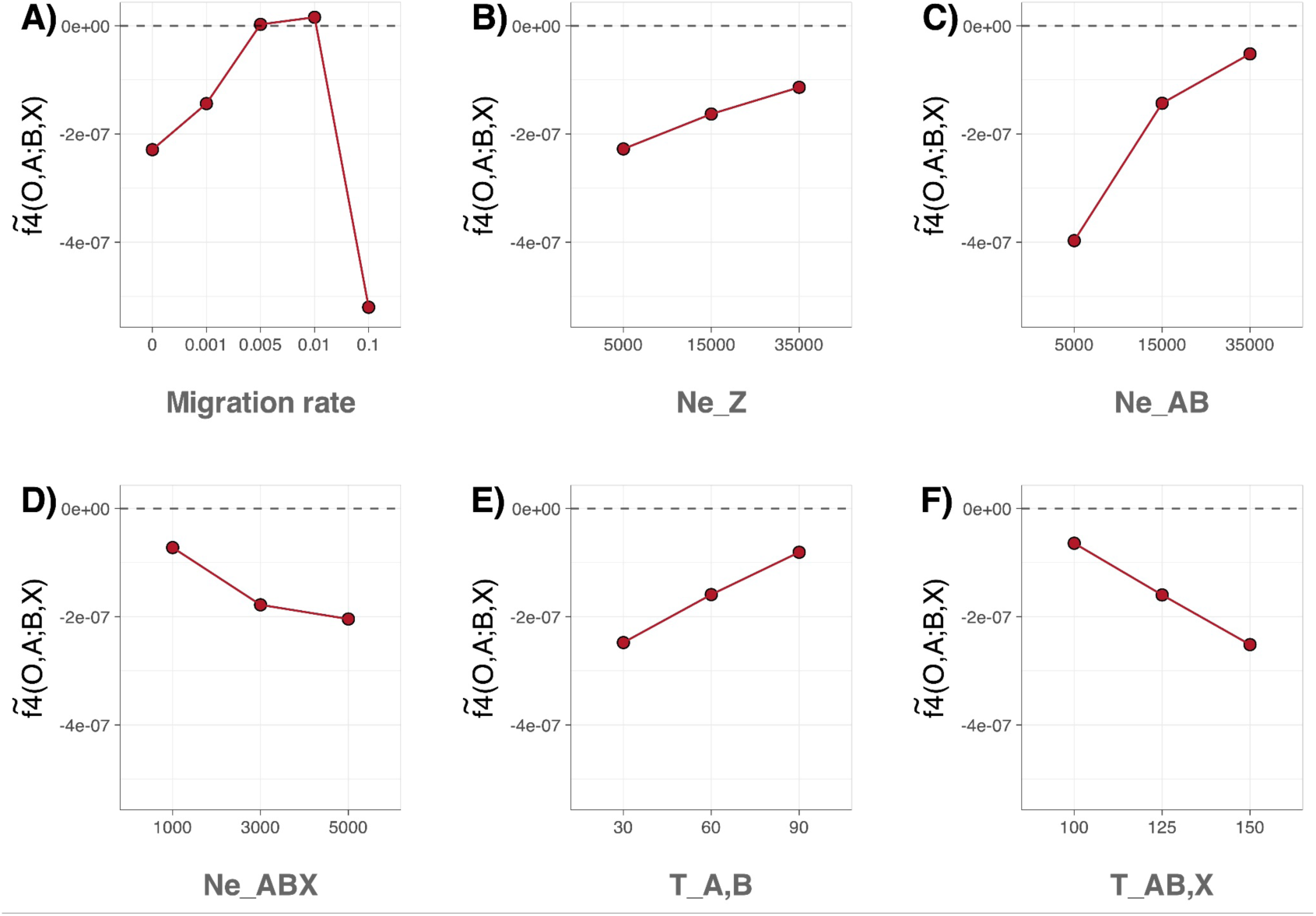
Change in *f_4_* statistics of the form *f_4_(O, A; B, X)* in population genetic simulations, where all parameters except the one in question were fixed at their median values. The median *f_4_* value for all replicates is plotted per variable.

As opposed to results involving low-level migration (m=0.001 to m=0.01), at m=0.1, *f_4_(O, A; B, X)* values shift back to negative values, i.e. sister attraction (Figure 4,5, S4). This happens because high migration rates from Z lead to the homogenization of A and B with Z. In other words, A and B become part of the Z gene pool. The drop in *f_4_* values is particularly strong if Ne_Z_ is low: e.g. at m=0.1 and Ne_Z_ = 5k all results are negative (Figure 4,5, S4). The theoretical model in Figure 3 also predicts a similar behaviour when migration probabilities are high.

We then asked whether, in simulations where we detect sister repulsion between A and B in *f_4_*-statistics, we would still observe A and B clustering separately from X in the MDS analysis, a pattern we had observed in the empirical data (Figure 1). For this, we chose one random case from each scenario and calculated outgroup-*f_3_* statistics. We then compared the mean distance between X to A and B [<u>*d*</u>_mds_(X,A/B)] versus the distance between A and B [*d*_mds_(A,B)] in MDS space. In 99% of all cases, populations A and B clustered separately from X, i.e. *d*_mds_(A,B) was lower. This included scenarios where 100% of the *f_4_* values signified sister repulsion (Figure 6A,B). Meanwhile, under low-level migration we also observed *f_3_(O; A, B) < f_3_(O; A/B, X)*, reflecting sister repulsion (Figure 6C). Hence, similar to our observation with the East Mediterranean Bronze Age genomes (Figure 1B,D), MDS-based clusterings in our simulations contradict pairwise *f_3_* statistics.

**Figure 6:**
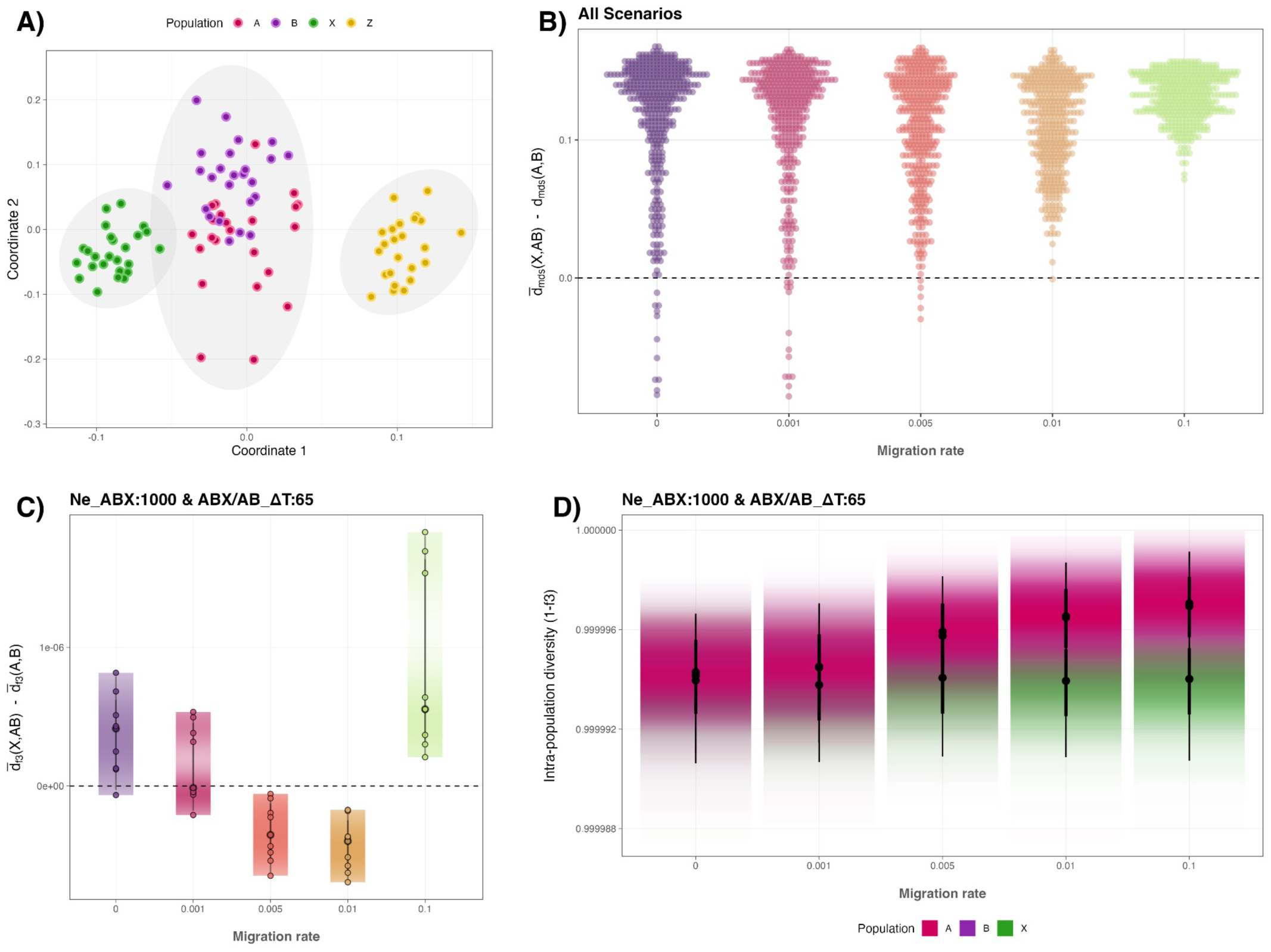
**(A)** MDS of simulated populations A, B, X and Z, using 1 - outgroup-*f_3_* values as genetic distances, for one sample scenario where m_Z->A.B_ : 0.01,Ne_Z_ : 15000, Ne_AB_ : 15000, Ne_ABX_ : 1000, and ΔT_X-AB,A-B_ : 65. **(B)** MDS clusterings between A and B versus X and A,B, calculated by the Euclidean distances between the mean MDS coordinates of the corresponding populations, for all scenarios. <u>*d*</u>_mds_(X,AB) is the mean distance of X to A and B. d_mds_(A,B) is the distance between A and B. Positive values for <u>*d*</u>_mds_(X,AB) - d_mds_(A,B) represent higher distance of X to A and B, thus distinct clustering of X from A and B. **(C)** Change in f3 values with migration rate from Z for scenarios where Ne_ABX_ : 1000 and ΔT_X-AB,A-B_ : 65. <u>*d*</u>_f3_(X,A/B) is the mean 1-f3 distance of X to A or to B. <u>*d*</u>_f3_(A,B) is the mean 1-f_3_ distance between A and B. Negative values for <u>*d*</u>_f3_(X,A/B) - <u>*d*</u>_f3_(A,B) show higher affinity between A or B and X than between A and B. **(D)** Intra-population diversity differences between populations A,B and X in relation to migration rate from Z, for scenarios where Ne_ABX_ : 1000 and ΔT_X-AB,A-B_ : 65. The diversities are measured by pairwise 1-outgroup f_3_ values per population. At migration rates 0 and 0.001, the values for X have higher deviation and overlap with A and B, thus have lower visibility.

In these models, we might expect population differences in within-population diversity, which could be a source of information for inferring unknown admixture history. Indeed, in the Bronze Age dataset we had observed significantly higher diversity in genomes from Anatolia versus modern-day Greece (Figure 1C). In our simulations that involved gene flow from Z, we likewise observed elevated intra-population diversity in A and B versus X (Figure 6D).

We next studied sister repulsion under additional models and assumptions:

a. We asked how asymmetric Z-sourced gene flow, where only population B receives migrants from Z, would impact *f_4_* statistics. In this case, as might be expected, *f_4_(O, A; B, X)* tends to be positive, i.e. A shows sister repulsion. Conversely, *f_4_(O, B; A, X)* tends to be negative, i.e. B is still attracted to its sister A (Figure S5). This may also be anticipated from Figure 3: if *α*_B_=0, *F_4_* would always be negative.
b. We tested gene flow from Z into AB, the ancestor population of A and B. This yielded similar patterns to those observed with gene flow to A and B separately, in terms of both *f_4_*, MDS and diversity estimates (Figure S6).
c. Our main simulations comprise continuous gene flow from Z starting from the A-B split. We hence used single pulse migration models, and replicated our findings with continuous gene flow. As expected, these required higher levels of per generation gene flow to produce the same effect (Figure S7); to be compared with Figure 4).
d. Because the values used for the T_A-B_ parameter (30-90 generations) are higher than what might be anticipated for the split time of two East Anatolian populations, we tested shorter split times, of 5 and 10 generations. We could again replicate sister repulsion patterns whilst at higher rates of migration (Figure S8; to be compared with Figure 4).
e. We asked whether applying a more realistic setting for genomic data simulation would change the results. We repeated the simulations using the HapMap chromosome 1 recombination map (The International HapMap Consortium) and found that the results remained the same (Figure S9).

These results reveal how low-level gene flow from an external source (Z) can lead to sister repulsion, i.e. closely related pairs of populations (A and B) showing higher affinity to a lineage that has split earlier from them (X) in *f_3_* and *f_4_* statistics. This effect happens because the f-statistics are point estimates that represent the average across diverse phylogenetic paths in a population’s demographic history. In our model, the effect depends on the distance of Z to XAB; therefore the introduction of so-called “deep ancestry” can readily shift the expected values. This also resembles the “ghost admixture” phenomenon when gene flow from distant sources can bias phylogenetic inference (Rogers and Bohlender 2015; Tricou *et al*. 2022).

Finally, we tested whether gene flow from X into A and into B independently (after the A-B split) might also lead to sister repulsion, without invoking the external source Z. We confirmed this was the case using both *f_3_*- and *f_4_*-statistics (Figure S10A). We further found that patterns observed both in the above simulations with external gene flow (Figure 4-6) and in the empirical data (Figure 1), namely lower diversity in X versus A/B and separate MDS clustering of X and A/B, can be replicated under gene flow from X, but only when the effective population size of X (Ne_X_) is low relative to Ne_A_ and Ne_B_ (Figure S10B,C).

Sister repulsion is an interesting case because in all the scenarios studied, A and B share more similar demographic histories with each other than with X, although this is not directly reflected in the *f_3_*- and *f_4_*-statistics. When sister repulsion is suspected, researchers may resort to alternative tools such as genetic clustering (e.g. MDS), qpAdm modelling (Harney *et al*. 2021) or using time stratifying variants (Speidel *et al*. 2024) to disentangle the signals. As a quick diagnostic of the problem, we highlight the method of randomly splitting one of the target populations, whereby if *f_4_* statistics show the repulsion signal between the two subgroups, sister repulsion is evident. Researchers may further be interested in evaluating whether post-split admixture from X or an external population Z is more likely, which may be evaluated using additional sources of information, such as local ancestry inference or archaeological/historical evidence for regional gene flow. In the empirical cases we covered, such as sister repulsion in Bronze Age East Anatolia, external gene flow from genetically distant populations (e.g. related to sub-Saharan Africa) may be a more parsimonious explanation than direct gene flow from the population being compared (e.g., Greece).

## Methods

### Ancient genomes

We used 79 published Bronze Age genomes from the East Mediterranean (Supplementary Table 1), from a publicly available dataset comprising the ancient individuals analyzed in Lazaridis et al., 2022, genotyped at the 1240K array positions (Lazaridis *et al*. 2022a). We only used Bronze Age genomes from East Anatolia and Greece produced with the 1240K enrichment method in order to avoid any technical bias that can arise when co-analyzing different data types, e.g. shotgun versus capture (Koptekin *et al*. 2023).

### Principal Components Analysis (PCA)

For the principal components analysis, we used the Human Origins Dataset (Lazaridis *et al*. 2014, 2016), containing 2,583 present-day genomes genotyped at the Affymetrix Human Origins Array positions, merged with the Bronze Age genomes. PCA was performed using *smartpca v.18140*, with option *lsqproject:YES*, implemented in *EIGENSOFT v.7.2.0* (Patterson *et al*. 2006). We projected the ancient individuals onto the first two principal components calculated from present-day Western Eurasian diversity based on populations from the Human Origins panel.

### *f_3_, f_4_* - statistics and Multidimensional scaling (MDS)

We calculated outgroup *f_3_-* and *f_4_*-statistics using *admixr v0.9.1,* which runs *AdmixTools v.7.0.2,* with default parameters (Patterson *et al*. 2012; Petr *et al*. 2019b). Within-group genetic diversities were estimated by computing pairwise 1 - outgroup-*f_3_*. All statistics were computed based on the 1240K dataset, using 3 modern-day Yoruba individuals as the outgroup. We estimated pairwise genetic distances with 1- outgroup *f_3_*, computed a distance matrix of 1-*f_3_* values, and performed a classical multidimensional scaling analysis using the *cmdscale* algorithm implemented in the R package *stats* (R Core Team, 2023).

The geodesic distances between sites for the comparison of genetic versus geographic distances were calculated using the R package *geosphere* (Hijmans 2010).

For the calculation of *f_4_*(Yoruba, Group1; Group2, Greece) statistics within the East Anatolian site Alalakh, we randomly split the 25 individuals in two groups of 20 & 5 and 15 & 10. Repeating each of the two groupings five times, we computed ten *f_4_* statistics in total.

### Simulations

We used *msprime v1.0* (Baumdicker *et al*. 2022) to simulate diploid sequences of 100Mb, where we sampled 25 individuals from each population. The divergence time of the population used as outgroup (ABXZO) was given as 200 generations ago, followed by the divergence of ABXZ after 20 generations. The effective population size of each population other than the ones given in Table 1 were fixed as 5k, i.e. populations ABXZO, O, ABXZ, X (except for scenarios where Ne_ABX_ = 1k and ΔT_X-AB,A-B_ = 65), A, and B. The mutation rate and the recombination rate were chosen as 1e-8 and 1.1e-8 per generation, respectively (Baumdicker *et al*. 2022). Each scenario had 100 replicates. One-way migration rates from Z were set the same into both A and B, except for one case (Ne_Z_, Ne_AB_, Ne_ABX_, ΔT_X-AB,A-B_) where we tested the effect of asymmetric migration by allowing gene flow (m=0.01) only into B. For scenarios where Ne_ABX_ = 1k and ΔT_X-AB,A-B_ = 65, one-way migration from X into A and B, migration from Z only into AB and single pulse migration models at times T_A-B_/2 and T_AB-X_/2 with additional migration rates (0.15 and 0.2) were also tested. All combinations of values for parameters m_Z->A.B_, Ne_Z_, Ne_AB_, Ne_ABX_, and ΔT_X-AB,A-B_ given in Table 1 were tested, in addition to scenarios where T_A-B_ was set to 5 and 10 generations ago. We also ran simulations using the HapMap recombination map for chromosome 1 (The International HapMap Consortium), under scenarios where Ne_ABX_ = 1k and ΔT_X-AB,A-B_ = 65. The outgroup-*f_3_* and *f_4_* statistics were calculated using *tskit v0.5.6*, which follows the calculation in Patterson et al., 2012, but the default *span_normalise=True* option we used normalizes the values by the total length of windows, which results in differences in the scale of the *f_3_* and *f_4_* values, *tskit* yielding smaller values than Admixtools. The approach for split-*f_4_* analysis explained above for real data was followed for a randomly chosen scenario where Ne_ABX_ = 1k, ΔT_X-AB,A-B_ = 65 and m=0.01. The *f_4_*(O, Group1; Group2,B) were calculated between groups of population A. The MDS analysis was performed the same way as above. In order to compare the MDS distances between populations, we used the mean MDS x and y coordinates of each individual, and calculated the mean coordinate point per population. We then calculated Euclidean distances between the mean coordinates of the corresponding populations.

## Data Availability

The code and parameters for generating the simulated data are deposited under https://github.com/goztag/msprime_fstats/tree/main.

## Acknowledgements

We thank Emilia Huerta-Sanchez, Torsten Günther, Benjamin Vernot, Pavlos Pavlidis, Dilek Koptekin, Yetkin Alıcı, Iosif Lazaridis and members of the METU CompEvo Team and the NEOMATRIX Collective for helpful discussions. The work was supported by the H2020 ERC Consolidator grant (no. 772390 NEOGENE to M.S.), H2020-WIDESPREAD-05-2020 TWINNING grant (no. 952317 NEOMATRIX to M.S.) and Scientific and Technological Research Council of Turkey (TÜBİTAK) through the 2210/A National Scholarship Programme for MSc Students (to G.A.).

## Notes

### Competing Interest Statement

The authors have declared no competing interest.

### Summary of Updates

Additional section on theoretical expectation of the effect and simulations with additional scenarios.

