## Supplemental Figures for "An explanation for the sister repulsion phenomenon in Patterson’s f-statistics"

### Supplementary Figures

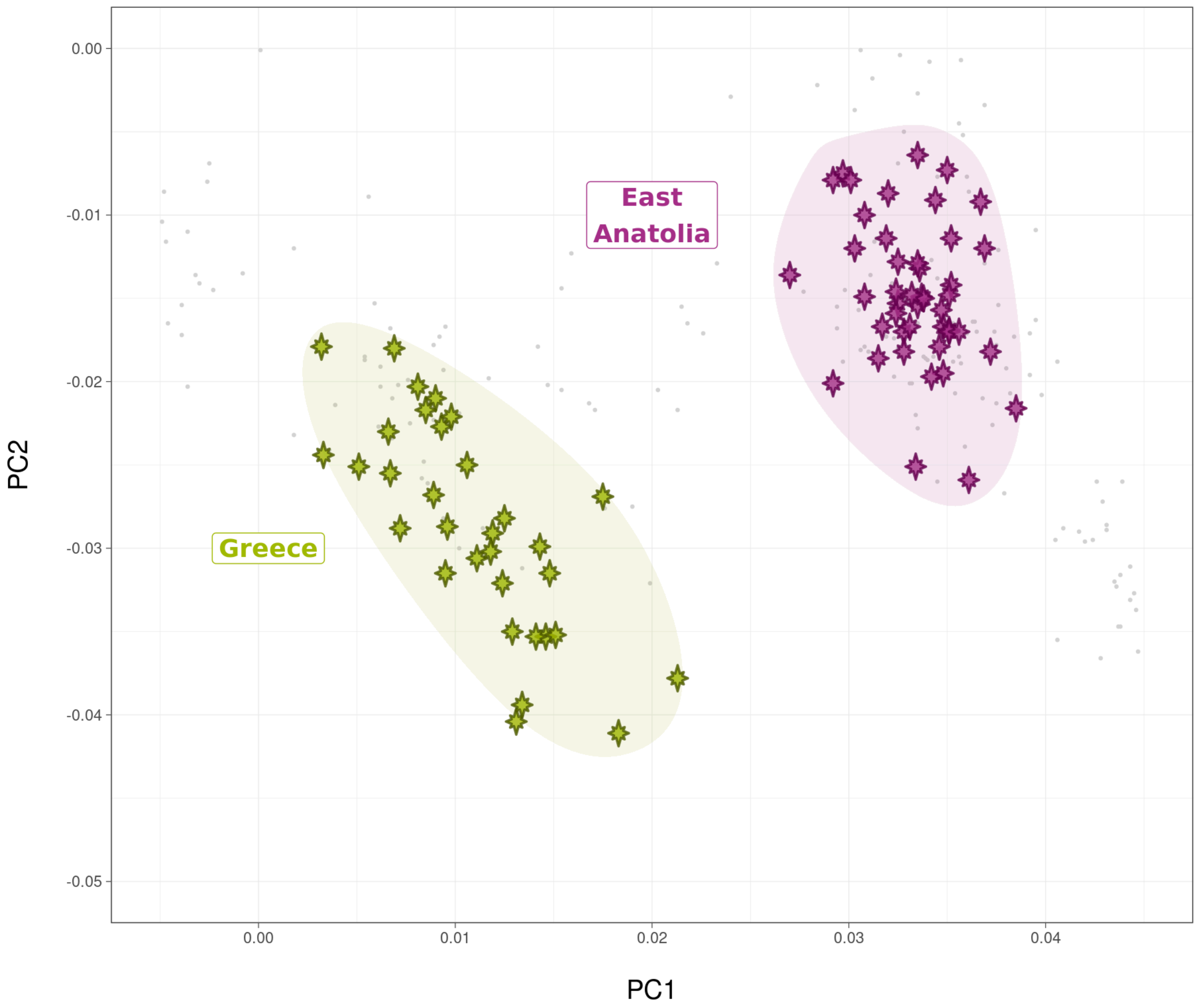


**Figure S1**. Principal components analysis (PCA) of the Bronze Age Greece and East Anatolian genomes, projected on the distribution of present-day genomes from the Human Origins panel (Patterson et al. 2012), indicated by gray points.

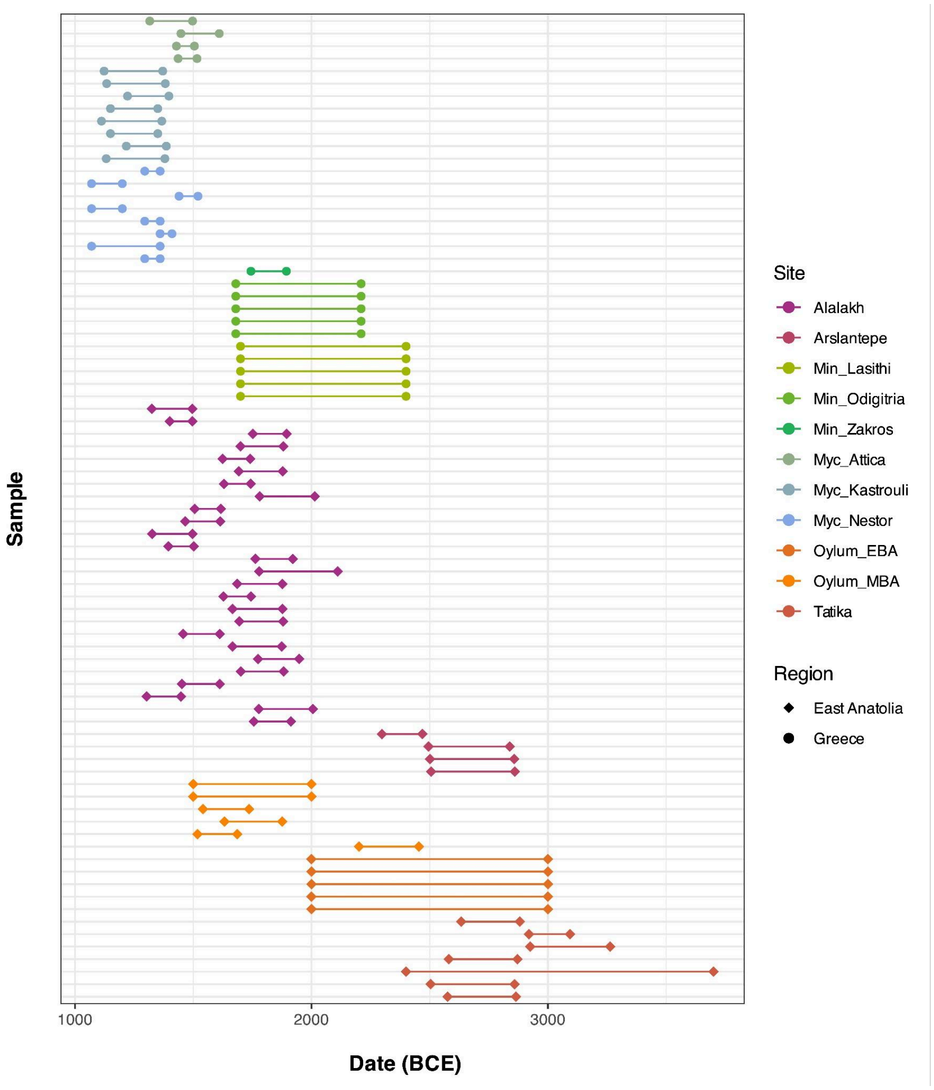


**Figure S2.** The ages of published genomes from Bronze Age Greece and East Anatolian analysed in this study. The age information was collected from the literature and are either C14- or archaeological context-based.

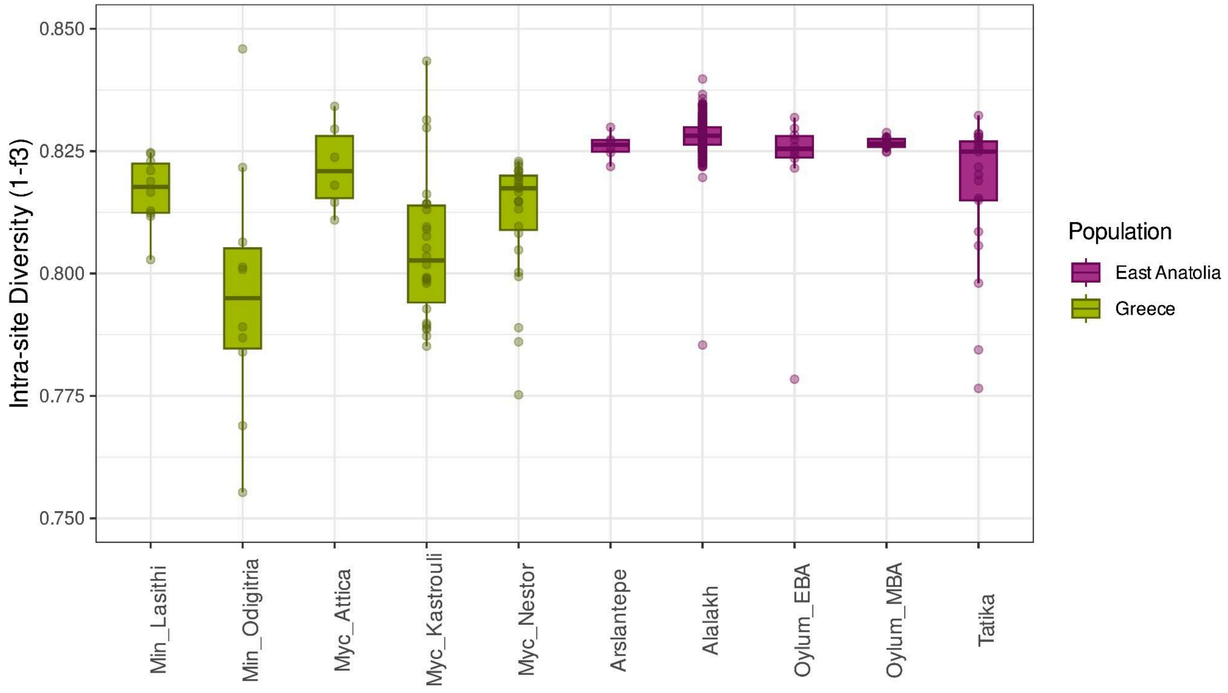


**Figure S3.** Genetic diversity estimates per site, calculated with 1-f3 statistics between all pairs of genomes from each site.

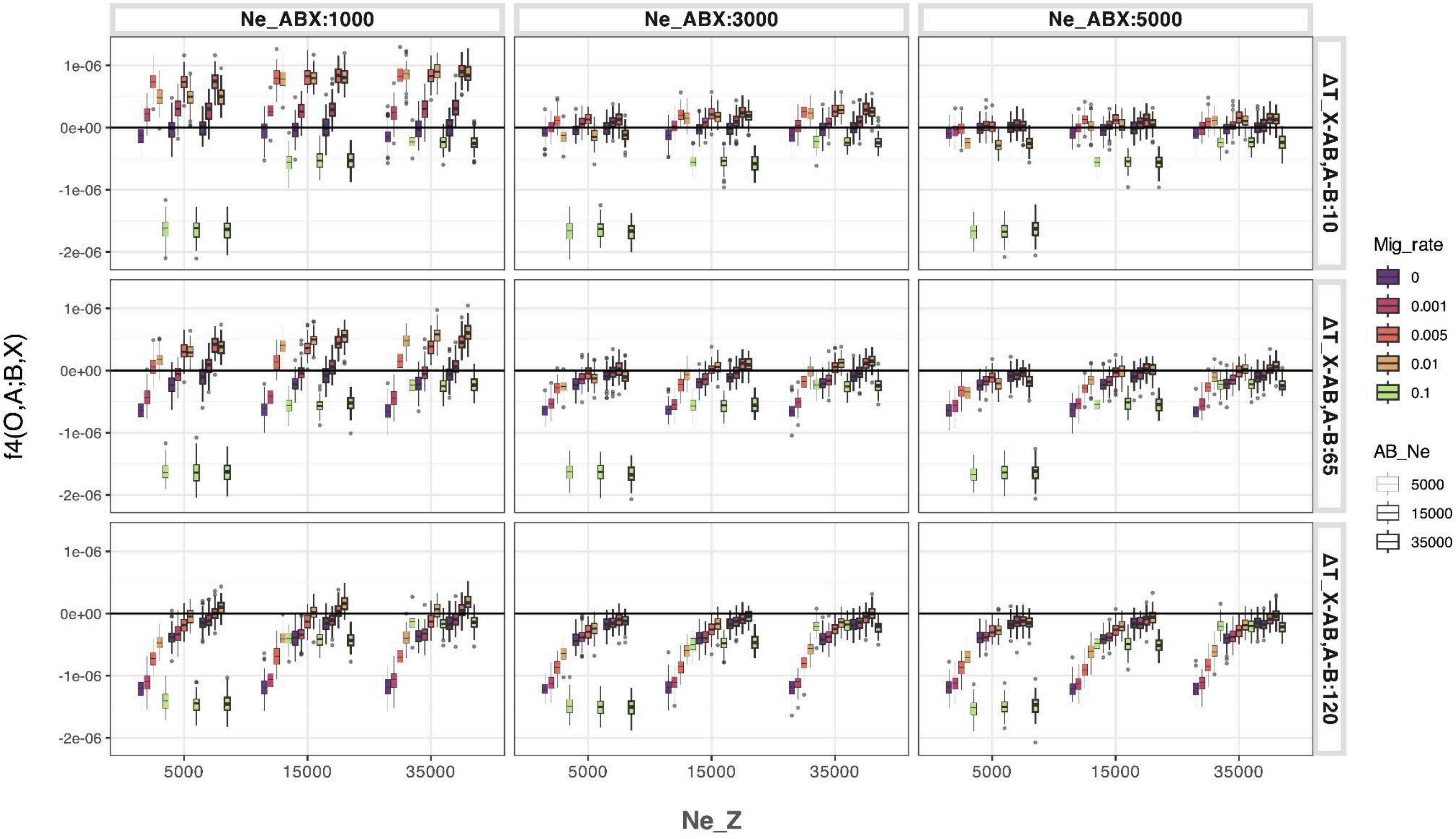


**Figure S4.** $f_4$ statistics of the form $f_4(O, A; B, X)$ depending on the varying parameters in Table 1. Negative $f_4$ values represent higher affinity between A and B (sister attraction), and positive values represent higher affinity between A and X (sister repulsion). Note that $f_4$ values in the first column and first two row panels, where $Ne_{ABX}$ = 1k, $Ne_Z$ > 5k, $\Delta T_{X\text{-}AB,A\text{-}B} \leq 65$ and $0.001 \leq m < 0.1$ are all positive.

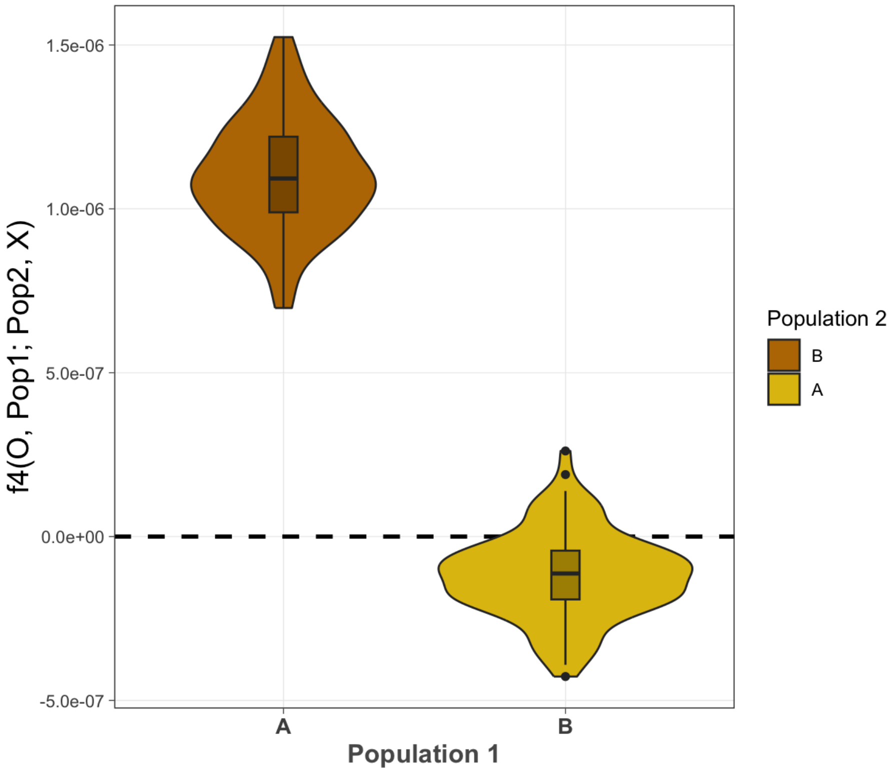


**Figure S5.** The effect of gene flow only into population B but not A. $f_4$ statistics of the form $f_4$*(Outgroup, Pop1; Pop2, X)*, where Pop1 and Pop2 are either population A or B, for the scenario where only population B receives external gene flow from Z. Positive $f_4$ values show higher affinity between X and Pop1 (sister repulsion), while negative $f_4$ values show higher affinity between Pop1 and Pop2 (sister attraction).

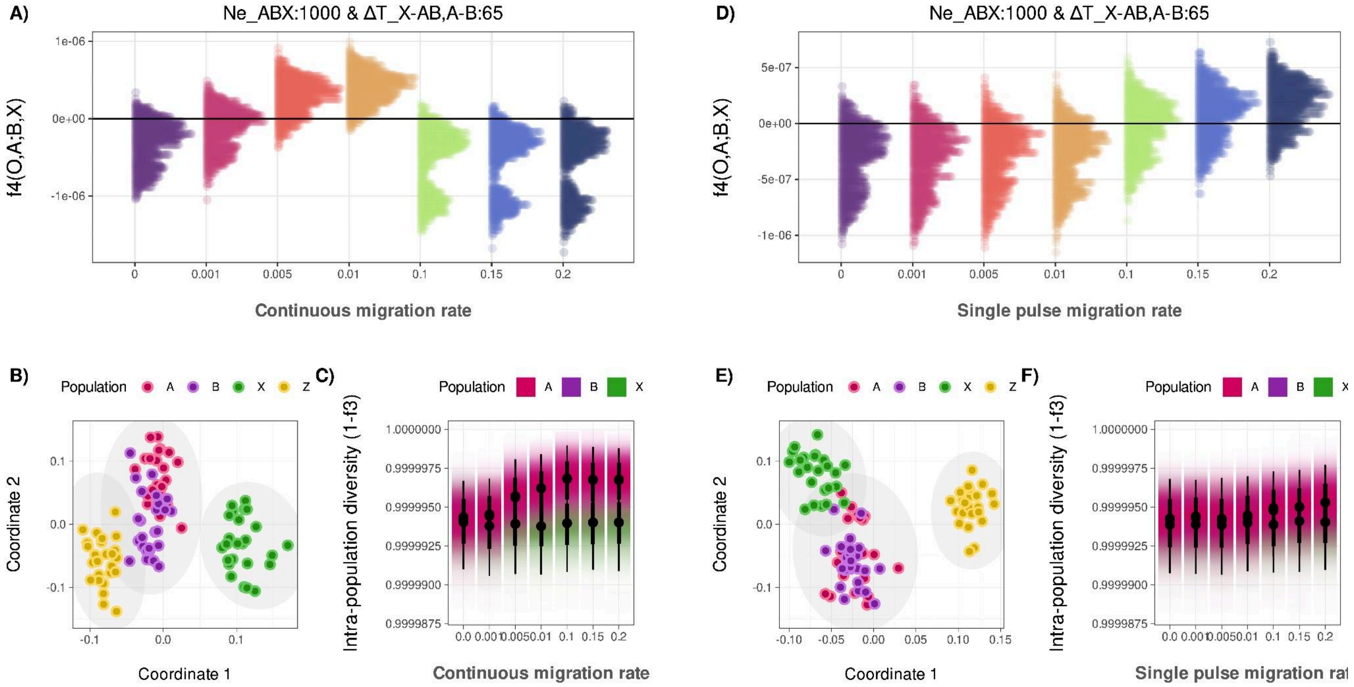


**Figure S6.** **(A)** $f_4$ statistics of the form $f_4(O, A; B, X)$ with continuous migration from Z to ancestral AB. **(B)** MDS summarizing pairwise 1 - outgroup $f_3$ distances between individuals from simulated populations where continuous $m_{Z\text{->}AB}$=0.01. **(C)** Intra-population diversity differences between populations A, B and X in relation to continuous $m_{Z\text{->}AB}$. At the values where X has higher deviation and overlaps with A and B, it has lower visibility. **(D)** $f_4$ statistics of the form $f_4(O, A; B, X)$ with single pulse $m_{Z\text{->}AB}$ at generation $T_{AB\text{-}X}$ / 2. **(E)** MDS summarizing pairwise 1 - outgroup $f_3$ distances between individuals from simulated populations where single pulse $m_{Z\text{->}AB}$=0.01. **(F)** Intra-population diversity differences between populations A, B and X in relation to single pulse $m_{Z\text{->}AB}$. At the values where X has higher deviation and overlaps with A and B, it has lower visibility.

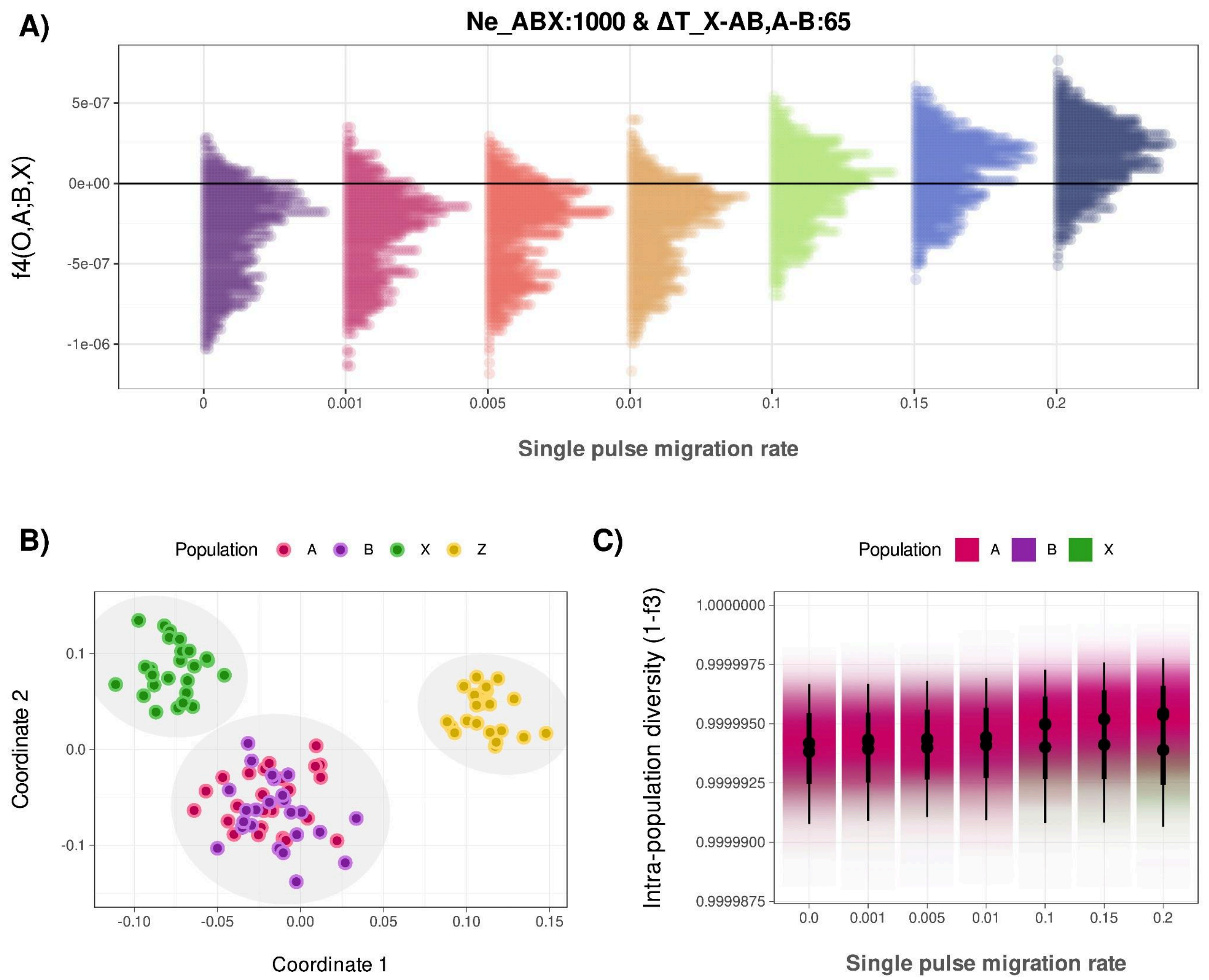


**Figure S7. (A)** $f_4$ statistics of the form $f_4(O, A; B, X)$ with single pulse $m_{Z\text{->}A,B}$ at generation $T_{A\text{-}B}$ / 2, for scenarios where $Ne_{ABX}$ : 1000, and $\Delta T_{X\text{-}ABX,A\text{-}B}$ : 65. Negative $f_4$ values represent higher affinity between A and B (sister attraction), and positive values higher affinity between A and X (sister repulsion). **(B)** MDS summarizing pairwise 1 - outgroup $f_3$ distances between individuals from simulated populations A, B, X and Z from scenarios where A and B receive gene flow from Z (m=0.01). **(C)** Intra-population diversity differences between populations A, B and X in relation to single pulse $m_{Z\text{->}A,B}$ , for scenarios where $Ne_{ABX}$ : 1000 and $\Delta T_{X\text{-}ABX,A\text{-}B}$ : 65. The diversities are measured by pairwise 1 - outgroup $f_3$ values per population. At the values where X has higher deviation and overlaps with A and B, it has lower visibility.

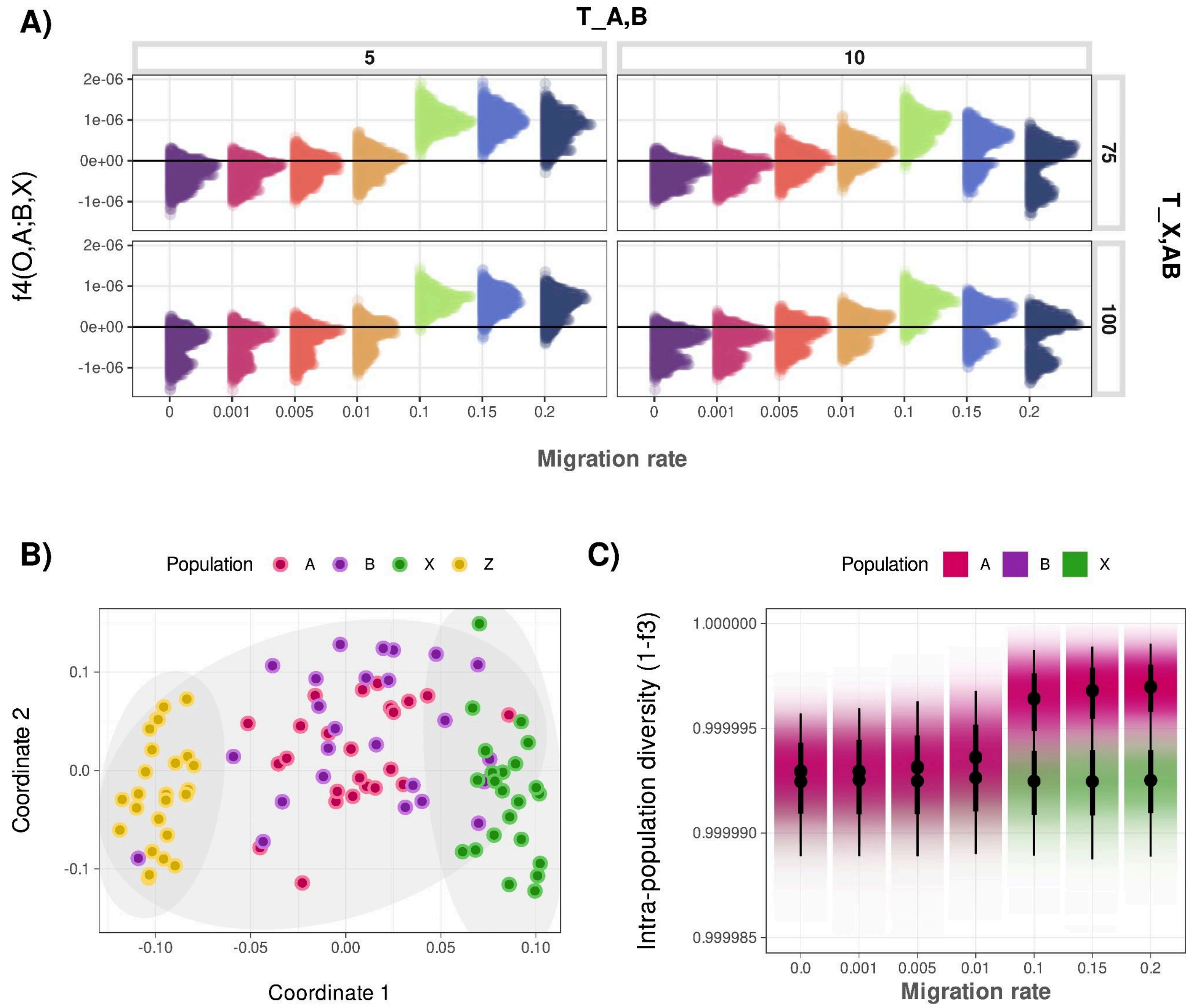


**Figure S8. (A)** $f_4$ statistics of the form $f_4(O, A; B, X)$ with $T_{A\text{-}B}$ and $m_{Z\text{->}A,B}$. Negative $f_4$ values represent higher affinity between A and B (sister attraction), and positive values higher affinity between A and X (sister repulsion). **(B)** MDS summarizing pairwise 1 - outgroup $f_3$ distances between individuals from simulated populations A, B, X and Z from scenarios where A and B receive gene flow from Z (m=0.1). **(C)** Intra-population diversity differences between populations A, B and X in relation to $m_{Z\text{->}A,B}$. The diversities are measured by pairwise 1 - outgroup $f_3$ values per population. At the values where X has higher deviation and overlaps with A and B, it has lower visibility.

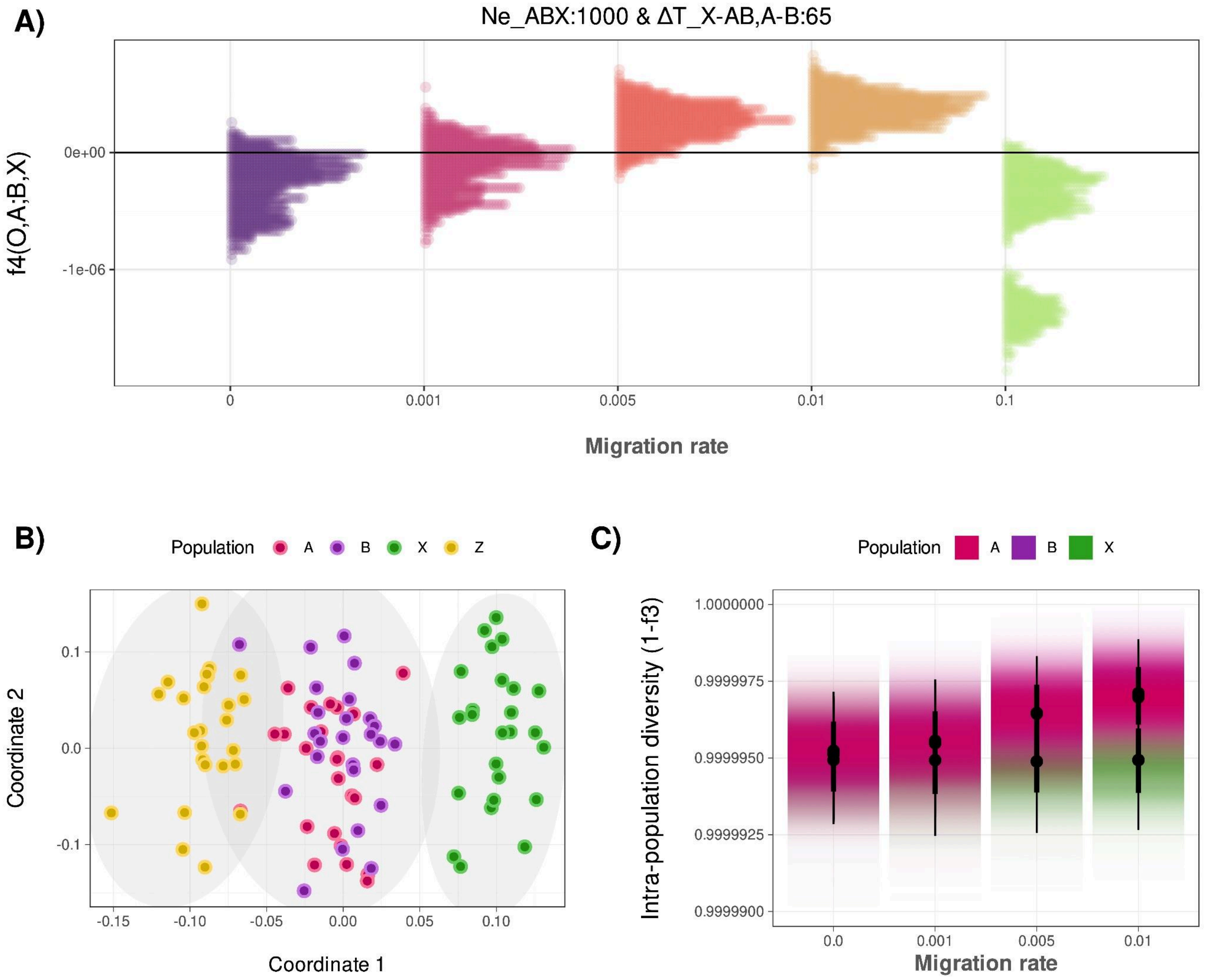


**Figure S9.** Simulation results based on HapMap chromosome 1 recombination map. **(A)** $f_4$ statistics of the form $f_4(O, A; B, X)$ depending on the varying $m_{Z\text{->}A,B}$, for scenarios where $Ne_{ABX}$ : 1000, and $\Delta T_{X\text{-}ABX,A\text{-}B}$ : 65. **(B)** MDS summarizing pairwise 1 - outgroup $f_3$ distances between individuals from simulated populations A, B, X and Z where $m_{Z\text{->}A,B}$=0.01. **(C)** Intra-population diversity differences between populations A, B and X in relation to $m_{Z>A,B}$. At the values where X has higher deviation and overlaps with A and B, it has lower visibility.

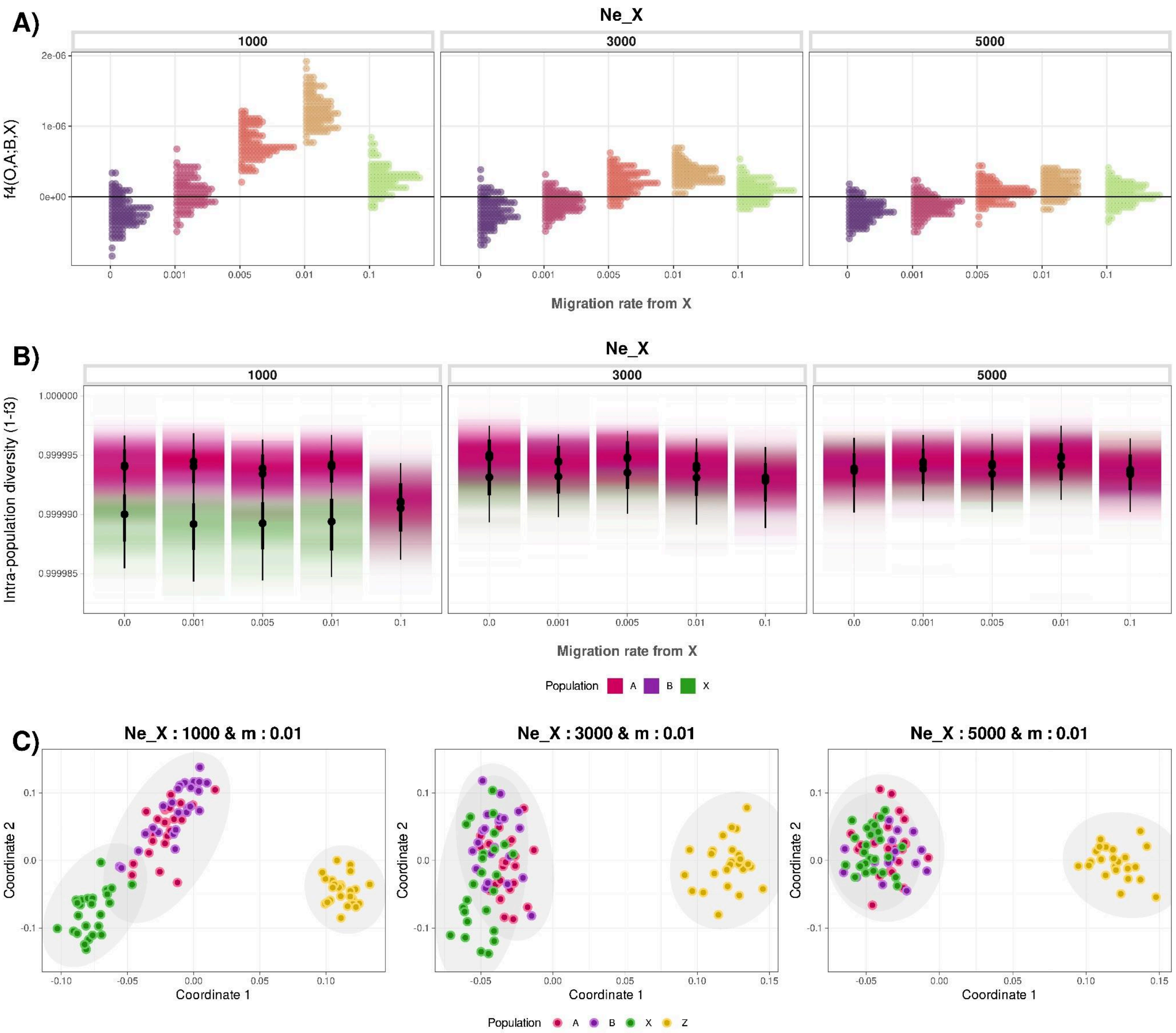


**Figure S10. (A)** $f_4$ statistics of the form $f_4(O, A; B, X)$ depending on the varying $Ne_X$ and $m_{X\text{->}A,B}$, for scenarios where $Ne_{ABX}$ : 1000, and $\Delta T_{X\text{-}ABX,A\text{-}B}$ : 65. Negative $f_4$ values represent higher affinity between A and B (sister attraction), and positive values higher affinity between A and X (sister repulsion). **(B)** Intra-population diversity differences between populations A, B and X in relation to $Ne_X$ and $m_{X\text{->}A,B}$ , for scenarios where $Ne_{ABX}$ : 1000 and $\Delta T_{X\text{-}ABX,A\text{-}B}$ : 65. The diversities are measured by pairwise 1 - outgroup $f_3$ values per population. At the values where X has higher deviation and overlaps with A and B, it has lower visibility. **(C)** MDS summarizing pairwise 1 - outgroup $f_3$ distances between individuals from simulated populations A, B, X and Z from scenarios where A and B receive gene flow from X (m=0.01), for different values (1k, 3k, 5k) of $Ne_X$.
